# Structural ubiquitin contributes to K48 linkage-specificity of the HECT ligase Tom1

**DOI:** 10.1101/2024.11.25.625210

**Authors:** Katrina Warner, Moritz Hunkeler, Kheewoong Baek, Anna Schmoker, Shourya S. Roy Burman, Daan Overwijn, Cyrus Jin, Katherine A. Donovan, Eric S. Fischer

**Author notes:** (E.S.F.).

## Abstract

Homologous to E6AP C-Terminus (HECT) ubiquitin ligases play key roles in essential pathways such as DNA repair, cell cycle control or protein quality control. Tom1 is one of five HECT ubiquitin E3 ligases encoded in the *S. cerevisiae* genome and prototypical for a ligase with pleiotropic functions such as ubiquitin chain amplification, orphan quality control and DNA damage response. Structures of full-length HECT ligases, including the Tom1 ortholog HUWE1, have been reported, but how domains beyond the conserved catalytic module contribute to catalysis remains largely elusive. Here, through cryogenic electron microscopy (cryo-EM) of Tom1 during an active ubiquitylation cycle, we demonstrate that the extended domain architecture directly contributes to activity. We identify a Tom1–ubiquitin architecture during ubiquitylation involving a non-canonical ubiquitin binding site in the solenoid shape of Tom1. We demonstrate that this ubiquitin binding site coordinates a structural ubiquitin contributing to the fidelity of K48 poly-ubiquitin chain assembly.

## Introduction

Ubiquitin is a 76 amino acid (aa) protein that acts as a post-translational modification to regulate a wide range of cellular processes in eukaryotes.^1,2^ These diverse roles are reflected in the plethora of mono- and poly-ubiquitylated adducts and substrates, as well as the diversity of ubiquitin linkage topologies.^3^ The process of ubiquitin transfer is tightly regulated by a class of enzymes called E3 ubiquitin ligases.^4^ Within this class, three major E3 ubiquitin ligase families exist: really interesting new gene (RING), ring-between-ring (RBR), and homologous to E6AP C-terminus (HECT).^5^ Dysregulation of several HECT ubiquitin ligases has been implicated in various cancer types, and neurological and neurodevelopmental disorders.^6^ To date, how the structural diversity of the non-catalytic modules present in this family direct enzymatic activity is an open question.

The HECT ubiquitin ligase family is characterized by a conserved ~350 aa HECT domain that folds into a bi-lobal catalytic module located at the C-terminus.^7,8^ During catalysis, ubiquitin forms a labile thioester intermediate with the HECT domain catalytic cysteine, forming an E3~Ub intermediate. The ubiquitin molecule is then transferred from the HECT catalytic cysteine directly to the substrate or onto another ubiquitin forming a ubiquitin chain. Structural intermediates representing the steps in this catalytic cycle have been visualized in the context of isolated HECT domains of various family members^9–14^ highlighting large scale domain rearrangements of the catalytic module that accompany ubiquitin transfer. Several recent studies have provided additional insights by visualizing full-length HECT ubiquitin ligases using cryogenic electron microscopy (cryo-EM), yielding the first structural views of full-length HUWE1,^15,16^ UBR5,^17–19^ and HACE1.^20^ Moreover, two recent studies captured HECT E3 ligases along K48-linked(UBR5, human)^19^ and K29-linked (Ufd4, *S. cerevisiae*)^21^ ubiquitin chain formation trajectories. These studies provide first snapshots into the role of domains beyond the HECT domain in facilitating ubiquitin transfer and in guiding chain specificity. However, it remains unclear whether other HECT ligases employ similar mechanisms, as HECT ligases beyond their HECT domains are structurally diverse. *S. cerevisiae* Tom1 and its human HUWE1 orthologs are essential HECT ubiquitin ligases, which act in a myriad of pathways associated with apoptosis, proliferation, orphan quality control, and differentiation.^22–30^ Human HUWE1 regulates a series of key stress response substrates, including Mcl1, p53, DDIT4, and Myc.^25,26,28,31^ Dysregulation of human HUWE1 is associated with X-linked intellectual disability and neurodevelopmental disorders, with several reported patient mutations in the ligase non-catalytic domains.^32,33^ Reconstituted ubiquitylation experiments demonstrate that the N-terminus contributes to ligase activity and ubiquitin loading,^16^ but the molecular mechanism of how these auxiliary domains contribute to activity remain elusive. Tom1, as the *<u>S. cerevisiae</u>* ortholog of HUWE1, regulates the same conserved pathways including orphan quality control, ubiquitin chain amplification, and DNA replication complex assembly.^34–36^ Tom1 lacks several disordered regions present in HUWE1, simplifying mechanistic characterization. In this study, we use Tom1, a functional and mechanistic ortholog of HUWE1, to dissect the molecular mechanism of ubiquitin transfer. We leverage direct visualization of a ubiquitylation reaction by cryo-EM to capture Tom1–E2–ubiquitin regulatory architectures that coordinate multiple ubiquitins. Using structure-guided mutations, we demonstrate that these ubiquitylation architectures are guided by the non-catalytic domains to contribute to faithful formation of K48 poly-ubiquitin chains.

## Results

### Tom1 shares key activities with HUWE1

First, we investigated whether HUWE1 and Tom1 are functional orthologs. We leveraged a previously reported addback model of Tom1, in which deletion of Tom1 in budding yeast results in a temperature sensitive phenotype.^34^ Adding back catalytically active Tom1 complemented the growth deficiency observed at elevated temperatures but introducing a catalytically inactive mutant (C3235S) was unable to rescue growth (**Figure 1A**), indicating the dependence on Tom1’s catalytic activity. Similarly, adding back catalytically active HUWE1 was able to rescue the growth deficiency, but not the catalytically inactive C4341S mutant (**Figure 1A**). Together, these results suggest that HUWE1 preserves the catalytic residue of Tom1 in this temperature sensitive phenotype.

**Figure 1.**
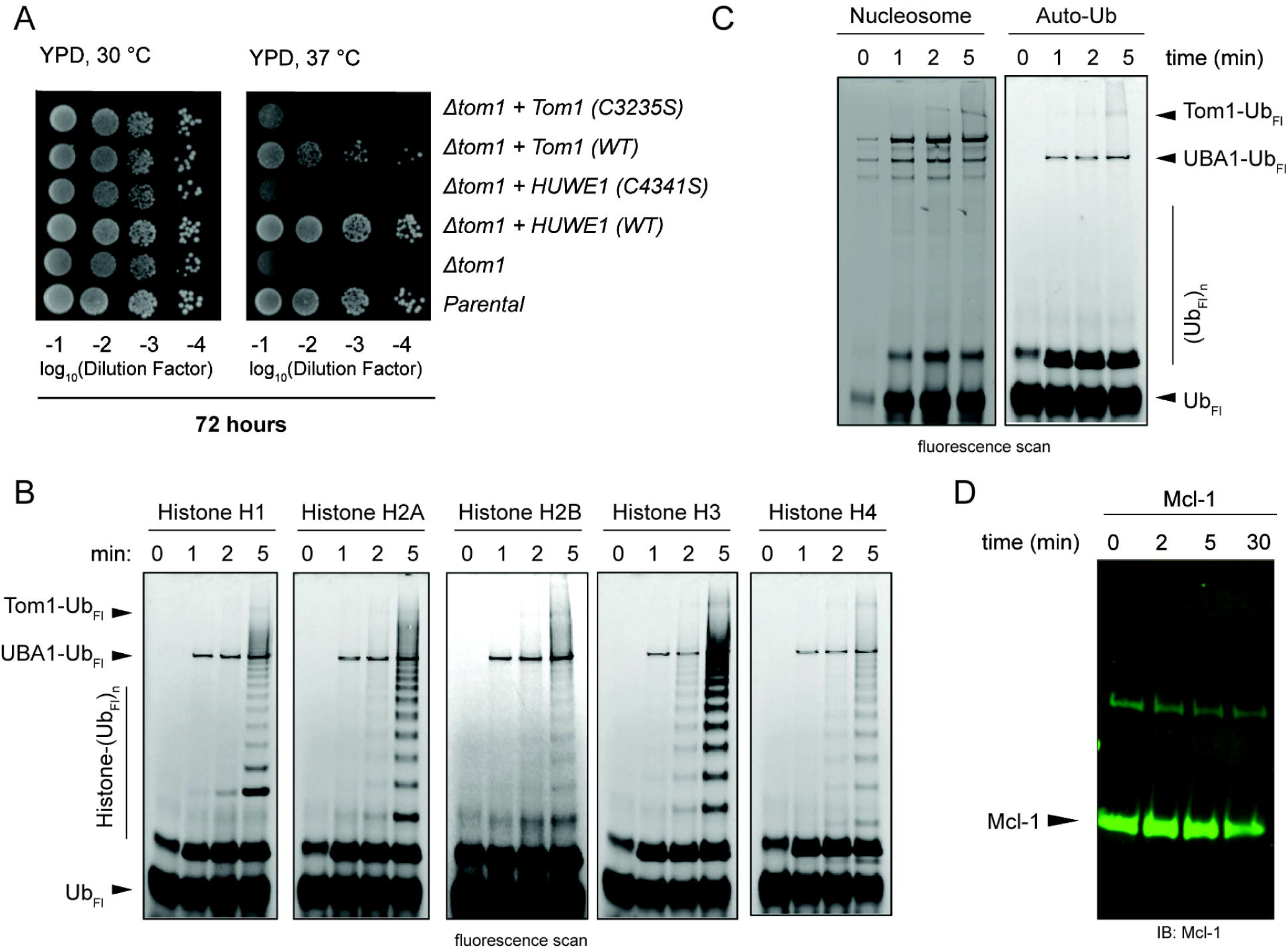
Tom1 shares key activities with HUWE1. **(A)** Cells of the indicated genotypes are spot plated on YPD and incubated at 30°C or 37°C for three days. **(B)** Ubiquitylation assay. Fluorescent ubiquitin transfer resulting from in vitro ubiquitylation of orphan histone substrates visualized on SDS-PAGE gel. **(C)** Ubiquitylation assay. Fluorescent ubiquitin transfer resulting from in vitro ubiquitylation of native nucleosomes or auto-ubiquitylation visualized on SDS-PAGE gel. **(D)** Ubiquitylation assay. Western blots using the indicated antibodies establishing Mcl-1 is not an *in vitro* substrate of Tom1.

We sought to understand whether functional and structural conservation extends to substrate recognition and ubiquitylation. We characterized the biochemical activity of Tom1 in the context of a cellular pathway shared among orthologs. Tom1, HUWE1 and HUWE1_N_ are known to function in orphan histone quality control.^15,23,34,37^ Using *in vitro* reconstitution and ubiquitylation assays, we demonstrated that while Tom1 ubiquitylates a series of orphan histones *in vitro*, it is inactive against recombinant nucleosomes (**Figures 1B, 1C**), highlighting the role of Tom1 in the context of orphan quality control. Tom1, akin to its *Nematodica spERT5* ortholog HUWE1_N_, does not include the BH3 domain that mediates Mcl-1 recognition in human HUWE1.^15,16,28^ Indeed, Tom1 showed no activity against Mcl-1 (**Figure 1D**), thus highlighting the specificity of substrate recognition mediated by the BH3 domain. Taken together, these results demonstrate that although HUWE1 evolved to have additional substrates, the core substrates and activities remain shared by Tom1 and HUWE1.

### Structure of Tom1 reveals conserved ring architecture

The Tom1 domain architecture is tightly conserved among its orthologs (**Figure 2A, S1B**).^15,16^ To establish this conservation at the structural level as well, we determined the structure of full-length Tom1 using cryo-EM (**Figure 2B**). Two-dimensional (2D) class averages revealed that Tom1 is conformationally heterogeneous (**Figure S1C**), similar to previous observations with HUWE1.^15,16^ Further classification demonstrated that the ligase engages in a dynamic motion where either the N- and C-terminal domains interact to form a “closed-ring” architecture, or the N-terminus moves away from the C-terminus to form a dynamic “open” conformation (**Figure S2B, S3C**). To minimize this conformational heterogeneity and obtain a high-resolution structure, we crosslinked Tom1 with BS3 to capture it in the closed conformation (**Figure S1C, Movie S1**). After several rounds of 3D classification and refinement, we obtained a 3D reconstruction that was refined to an overall resolution of ~3.0 Å. To ensure that BS3 crosslinking did not introduce artifacts, we collected a second dataset without crosslinking and obtained a similar reconstruction, yet at a lower overall resolution of ~3.8 Å (**Figure S2B**). Since the resulting maps were nearly identical, we built a high resolution model into the higher quality map from the BS3-crosslinked sample (**Figure 2C**).

**Figure 2.**
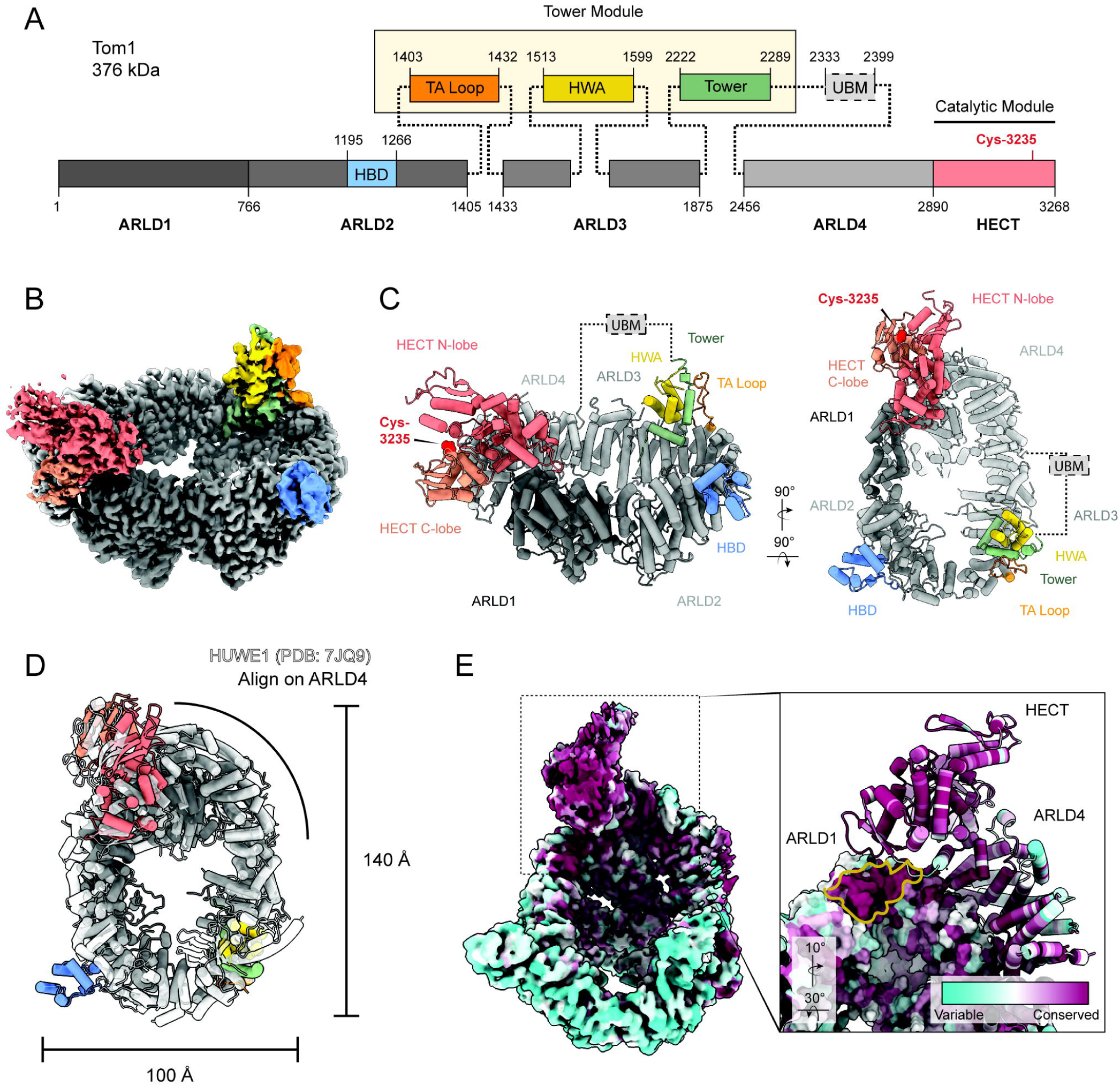
Structural characterization of full-length Tom1. **(A)** Domain map of full-length Tom1, indicating domain boundaries and amino acids stretched by ARLD1-4. Color scheme for the HECT and ARLD interfaces are kept constant throughout the remaining figures. **(B)** Consensus cryo-EM map of full-length Tom1, with domains colored as in Figure 2A. **(C)** Cartoon model of Tom1 in two orientations, with domains labelled. The UBM domain is not visible in the reconstructions. **(D)** Alignment of Tom1 and HUWE1 (PDB: 7JQ9) reveals the dimensions of the closed-ring architecture is tightly conserved. **(E) (Left)** Consensus cryo-EM map of full-length Tom1 in two orientations, colored to depict conservation analysis among 125 Tom1 orthologs. Highly conserved residues are depicted in maroon and variable residues are depicted in cyan, as determined by ConSurf.^57–60^ **(Right)** Rotated, magnified view of the extended ARLD1-ARLD4 contact interface formed in the closed-ring architecture. Surface representation of ARLD1 is displayed. A highly conserved patch proximal to the HECT domain and ARLD1–ARLD4 ring closure is outlined in yellow.

Tom1 forms an alpha solenoid assembly of near identical dimension to human HUWE1^16^ measuring ~140 Å × 100 Å × 100 Å (**Figure 2D**). The Tom1 central ring architecture comprises four armadillo repeat-like domains: ARLD1 (aa 1–750), ARLD2 (aa 766–1405), ARLD3 (aa 1433–1875), and ARLD4 (aa 2456–2889) (**Figure 2B, 2C**). This domain architecture is conserved among reported structures of full-length HUWE1 and HUWE1_N_ ^15,16^ (**Figure 2A, S1B**). The helical repeats in ARLD1 and HECT domain of Tom1 and HUWE1 align tightly domain-wise with root-mean-squared-deviations (RMSDs) of 2.5 and 1.4 Å, respectively (**Figure S1D**). The catalytic module (aa 2890– 3268) is nestled above the plane of the ring with the catalytic cysteine (C3235) vectored perpendicular to the closed-ring assembly (**Figure 2C, S1D**). The closed-ring architecture is stabilized through a 1,122 Å^2^ solvent-accessible surface area buried between ARLD1 and ARLD4 (**Figure 2C**). Conservation analysis of Tom1 revealed strong conservation at the ARLD1–ARLD4 interface, the HECT catalytic module, and an N-terminal patch close to the HECT and ARLD-ring closure (**Figure 2E, S1E**). Though the WWE domain present in HUWE1 is absent in Tom1, the helix-turn-helix “Tower” (aa 2222–2289) is flexibly tethered to ARLD3 and anchored by a tower-associated “TA” loop (aa 1403–1432) as well as a HUWE1 WWE-associated motif (HWA) (aa 1513– 1599) within ARLD2 (**Figure 2C, S1B**). A UBM module (aa 2333–2399), unresolved in our map, is tethered above the plane of the ring and anchored by ARLD3 and the “Tower” motif (**Figure 2C**). A previously uncharacterized four-helix bundle domain (HBD) (aa 1195–1266) protrudes out from ARLD2 and is unique to Tom1 (highlighted in blue in **Figure 2C**). The UBM, Tower, and HWA are highly (**Figure S1E**), and there are profound structural similarities among the three orthologs. Importantly, Tom1 lacks several flexible domains present in human HUWE1 (**Figure S1B**), facilitating further structural characterization. Overall, the structural and functional similarity between Tom1, HUWE1_N_, and HUWE1 helps explain the conserved core roles between orthologs.

### Conformational landscape of Tom1 during active ubiquitylation

To investigate the structural landscape during Tom1 mediated ubiquitylation, and whether the non-catalytic domains are involved in this process, we plunge-froze Tom1 while ubiquitylating H2B and visualized via cryo-EM. We separated the ubiquitylation reaction into two pre-incubated mixtures, one containing Tom1 and histone H2B substrate, and the second containing a “pre-charge mixture” consisting of UBA1, UBE2D2, Mg-ATP, and ubiquitin (**Figure 3A**). The reaction was initiated by adding the two mixtures and monitored via SDS-PAGE on a matched sample using fluorescently labelled H2B by BODIPY-maleimide (BODIPY-H2B) to visualize substrate ubiquitylation (**Figure 3B**). The samples were plunge-frozen 90 seconds post reaction initiation to capture the early steps of this reaction. Heterogeneous refinement of the resulting dataset produced three dominant Tom1 conformations: ligase open (**Figure 3B**, similar to Figure S2C), ligase closed (**Figure 3B**, similar to Figure 1A, S3B), and a complex that harbored Tom1 together with ubiquitin and E2 (**Figure 3B**). Expanding this complex that harbors Tom1-E2-UB showed different conformations with varying occupancy of the ubiquitin or E2 (**Figure 3C**): Tom1 associated with (class i) a mono-ubiquitin coordinated between ARLD1 and the HECT domain; (class ii) both a mono-ubiquitin bound ARLD1 and di-ubiquitin bound to the catalytic HECT N-lobe; (class iii) a mono-ubiquitin coordinated by ARLD1, two ubiquitin molecules bound the HECT N-lobe, and an E2 bound the HECT domain; (class iv) a K48 di-ubiquitin coordinated by ARLD1, two ubiquitin molecules bound the HECT N-lobe, and an E2 bound the HECT domain; (class v) a K48 di-ubiquitin coordinated by ARLD1, along with an E2 between this K48 di-ubiquitin and ARLD1/2, and two ubiquitin molecules bound the HECT N-lobe; and (class vi) all E2 and ubiquitin binding modes (**Figure 3C, 3D**).

**Figure 3.**
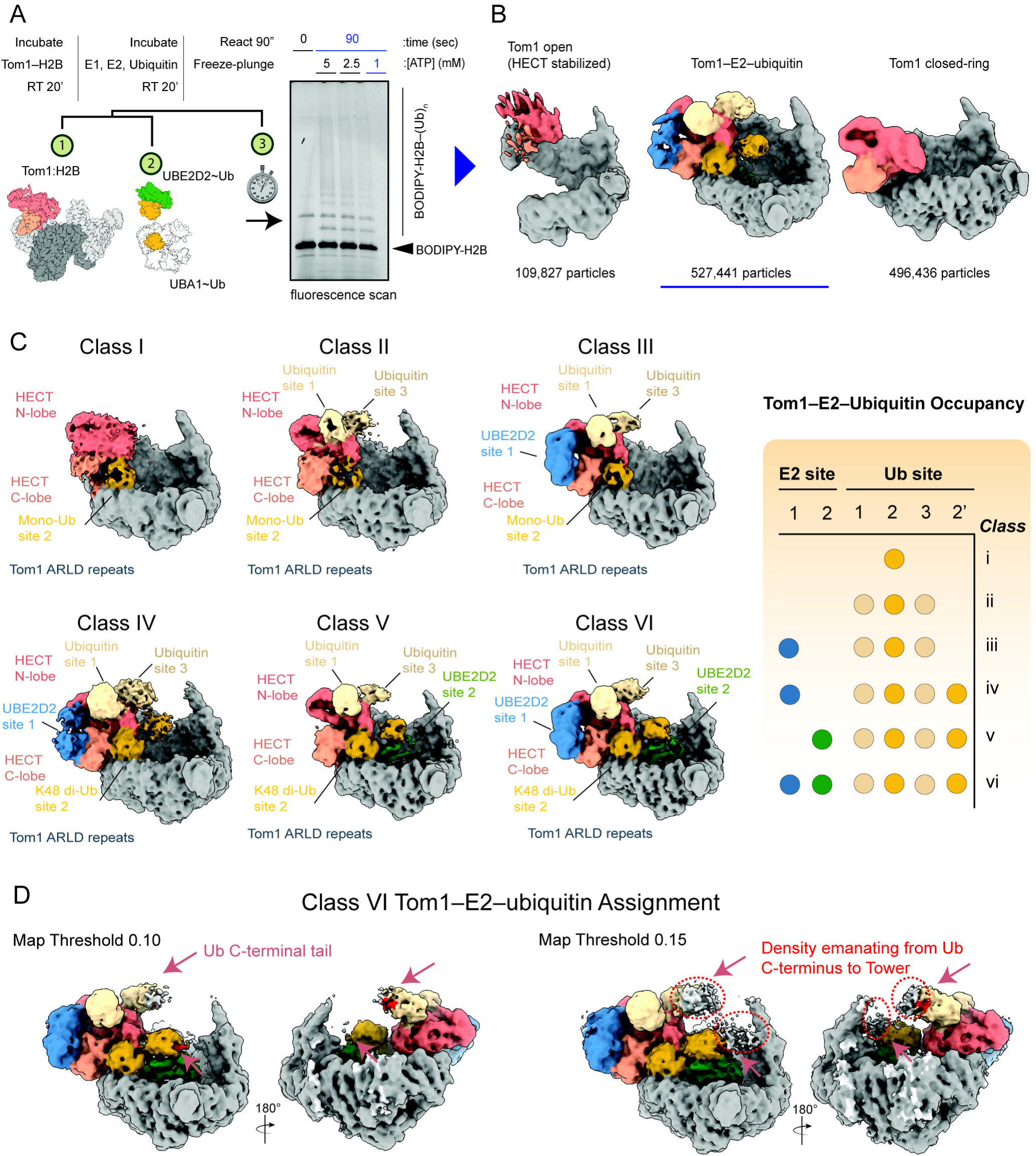
Dynamic E2–Ubiquitin occupancy during Tom1 ubiquitylation. **(A)** Schematic of the grid preparation from cryo-EM. (1) Tom1 and histone H2B were pre-incubated for 20 minutes at room temperature. (2) E1, E2 and ubiquitin were pre-incubated for 20 minutes at room temperature. (3) Each grid was double-blotted, once following a 45 second and subsequently following a 90 second co-incubation. (4) The reaction was plunge-froze in liquid ethane for downstream EM. Ubiquitylation assay performed under plunge-freeze conditions, depicting poly-ubiquitylation of covalently labelled histone H2B (H2B-BODIPY). **(B)** Landscape of Tom1 particles vitrified during in ubiquitylation cascade. Tom1 in the open conformation, the closed-ring conformation and a Tom1–E2–ubiquitin architecture are recovered. **(C) (Left)** Six particle clusters after 3D classification and heterogeneous refinement reveal dynamic occupancy of the various E2-ubiquitin binding sites along Tom1 during ubiquitylation. (Right) Graphical summary of E2 and ubiquitin occupancy classes determined by 3D classification. **(D)** Analysis of class 6 Tom1–E2–ubiquitin, depicted at two map thresholds (left 0.10, right 0.15). Density emanating from the C-terminal tail of either putative K48 di-ubiquitin (site 1 and 3 or site 2) extends towards the Tower domain.

### Tom1 N-terminal domain coordinates ubiquitin

To enhance our understanding of how the assembly of the maximally occupied (Class VI) Tom1–E2–ubiquitin state is structured, we processed a larger dataset at a higher magnification resulting in an overall global resolution of ~2.75 Å (**Figure 4A**). We identified a total of four ubiquitin molecules and two E2 enzymes bound to Tom1 as seen previously from the screening dataset (**Figure 4A**). Two ubiquitin molecules are bound to the HECT N-lobe (**Ub sites 1 and 3**), whose relative orientation is amenable to K48 linkage (**Figure 4A, 4B, S3V**) but the linkage itself could not be resolved. Ubiquitin site 1 was observed in previous studies of isolated HECT domain proteins^11,14^ (**Figure S3A**) and may coordinate substrate linked ubiquitin chains for amplification.^11^ In fact, we observed sparse density at low contour emanating from the C-terminus of the site 3 ubiquitin potentially directed towards the Tower domain which is presumably necessary for substrate engagement^15,38^ (**Figure 3D**), however the resolution is insufficient for confident assignment. A previously unreported K48 di-ubiquitin interaction (**Ub site 2**) is coordinated by the Tom1 N-terminal ARLD1 and HECT interface (**Figure 4C**). One molecule of UBE2D2 binds the canonical E2–HECT interaction site (**E2 site 1**) (**Figure 4B**),^9^ however, we observed no density for ubiquitin bound at the catalytic site of the E2 nor the HECT, presumably due to fleeting motion and the constant conjugation of ubiquitin to its next step. Surprisingly, a second UBE2D2 molecule was bound in a previously unreported site (**E2 site 2**) coordinated by Tom1 N-terminal ARLD1/ARLD2 along with the site 2 di-ubiquitin (**Figure 4D**). Taken together with the finding that the closed conformation is important for full catalytic activity of HUWE1,^15,16^ we demonstrated that the Tom1 HECT domain and ARLD1 jointly recognize K48 di-ubiquitin. Previously, it has been reported that back-binding of a structural ubiquitin to UBE2D2 increases ubiquitylation activity of UBE2D2.^39,40^ Here, we see a similar architecture of the site 2 di-ubiquitin back-binding to site 2 UBE2D2 (**Figure S3D**), which appears to be aiding the coordination of the bound K48 di-ubiquitin. The site 2 di-ubiquitin is coordinated by a 1,137 Å^2^ buried solvent accessible interface formed between N-terminal ARLD1 and the C-terminal HECT domain in the closed-ring conformation. To accommodate site 2 di-ubiquitin, the C-lobe rotates 35° towards the di-ubiquitin relative to the N-lobe forming the HECT–ubiquitin interface (**Figure S3B, Movie S2**). Moreover, the Tom1 ring expands by 22 Å. In addition, the N- and C-lobes of the HECT domain move towards the ubiquitin by 18 Å and 46 Å, respectively (**Figure S3C, Movie S2**). Previous studies of HUWE1 showed that the closed conformation is important for full catalytic activity.^16^ We demonstrate through Tom1 that the closed conformation coordinates K48 di-ubiquitin jointly by the HECT domain and ARLD1 through structural rearrangements, providing a potential rationale as to why the closed conformation is important for catalysis.

**Figure 4.**
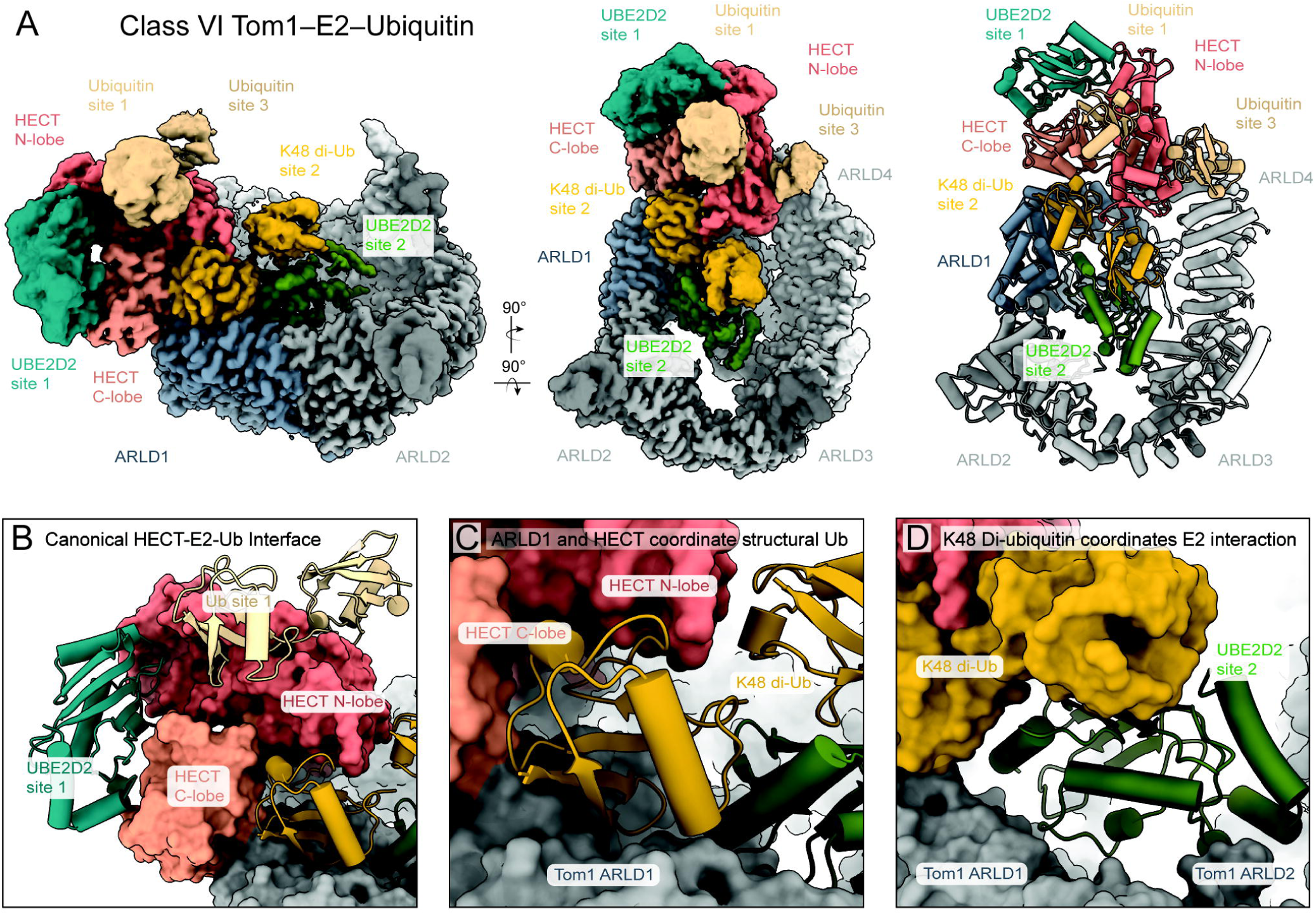
Structural analysis of Tom1–E2–ubiquitin binding. **(A)** Consensus cryo-EM map of full-length Tom1–E2–Ubiquitin complex in two orientations. Domains and bound proteins are labelled. Cartoon model of Tom1–E2– Ubiquitin complex at identical orientation. **(B)** Close-up view of site 1 E2 and site 1 ubiquitin binding mode. Site 1 UBE2D2 engages the HECT domain through a canonical E2–HECT interaction. Site 1 engages the HECT domain in a previously reported N-lobe exo-site. Tom1 is depicted as a surface representation. **(C)** Close-up view of site 2 K48 di-ubiquitin binding mode along Tom1 and site 2 UBE2D2. Tom1 and its domains are displayed in a surface representation. Site 2 UBE2D2 is colored in white. **(D)** Close-up view of site 2 E2–ubiquitin binding mode. Site 2 UBE2D2 engages one ubiquitin molecule through a literature reported ubiquitin back-binding interaction. Site 2 K48 di-ubiquitin and Tom1 are depicted as surface representations.

### Ubiquitin architecture regulates K48 chain selectivity

As the specificity of ubiquitin transfer relies on a precise geometry,^19,21,41^ we investigated how the observed ubiquitin coordinating architecture contributes to Tom1 activity and specificity. In Tom1, the highly conserved L615 residue in the N-terminal ARLD1 domain packs tightly against the hydrophobic F4/T12/I13/T14 patch on ubiquitin (**Figure 5A**). HECT C-lobe D3155 forms a salt bridge with S65 on ubiquitin (**Figure 5B**). We disrupted the N-terminal Tom1–ubiquitin packing interaction through the mutation of L615 to arginine (L615R), which increased overall substrate ubiquitylation relative to wild type (WT) (**Figure 5C**). Similarly, the HECT C-lobe mutation D3155R increased activity relative to WT. Tom1 and its orthologs are reported to selectively form and amplify K48 poly-ubiquitin chains.^15,35^ Therefore, we asked whether disrupting this E2– ubiquitin architecture influenced Tom1-mediated chain specificity. Quantitative mass spectrometry revealed that the Tom1 site 2 di-ubiquitin disrupting mutant, L615R, reduced Tom1 selectivity for K48 chain formation (**Figure 5D**). Similarly, detection of K48 ubiquitin linkages by immunoblot was diminished in Tom1 (L615R) relative to Tom1 (WT) in ubiquitylation assays (**Figure 5E**). Collectively, these results show that the site 2 ubiquitin contributes to K48 ubiquitin linkage specificity of Tom1.

**Figure 5.**
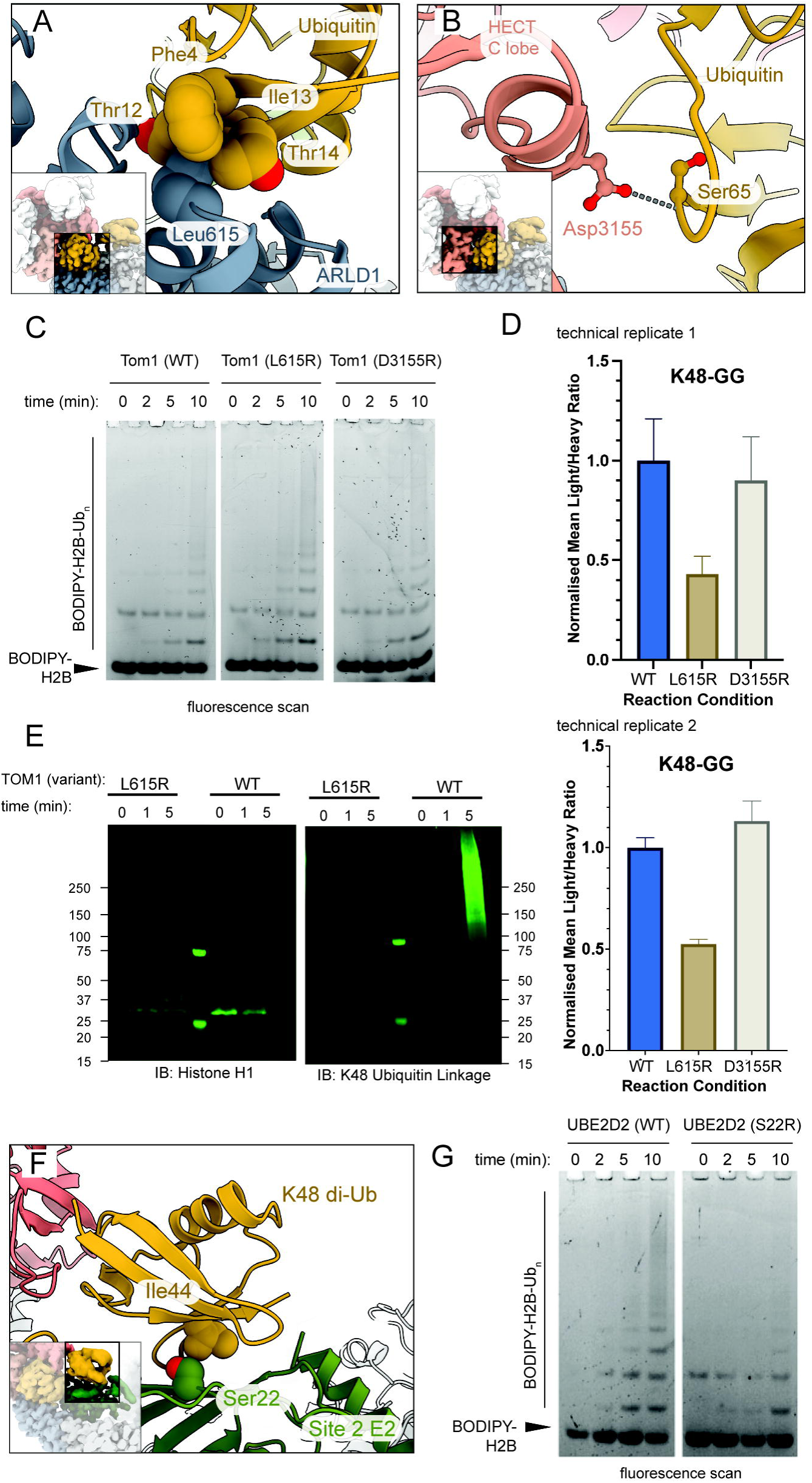
Structural ubiquitin regulates ligase activity and chain specificity. **(A)** Close-up view of Tom1 ARLD1–Ubiquitin site 2 contacts. Tom1 L615 is buried in a hydrophobic patch on ubiquitin. **(B)** Close-up view of Tom1 HECT C-lobe–Ubiquitin site 2 contacts. Tom1 D3155 forms a salt bridge with ubiquitin site 2 S65. **(C)** Ubiquitylation assay. Fluorescent H2B~BODIPY visualized on SDS-PAGE gel, resulting from in vitro ubiquitin of reaction. **(D)** Mass-spectrometry of in vitro histone H2B ubiquitylation assay. Depicted mean normalized light-to-heavy ratio of K48 di-Gly peptide relative to C-terminal Ubiquitin peptide (loading control) by Tom1 mutant reacted. Data represented as mean ± SEM (n=3). Two technical replicates depicted. **(E)** Ubiquitylation assay. Western blots using the indicated antibodies characterizing WT vs L615R mutant activity against histone H1. **(F)** Close-up view of UBE2D2–Ubiquitin back-binding contacts. UBE2D2 S22 forms a hydrophobic packing interaction with site 2 di-ubiquitin I44. **(G)** Ubiquitylation assay. Fluorescent H2B~BODIPY visualized on SDS-PAGE gel, resulting from in vitro ubiquitin of reaction.

E2 back-side interactions have been reported to stimulate ubiquitin transfer.^4,39,42–49^ Given the position of site 2 UBE2D2 relative to the ARLD-repeats, we questioned how disruption of the back-side interactions would influence activity. UBE2D2 at site 2 back-binds ubiquitin’s I44 hydrophobic patch (**Figure 5F, S4D**). Disruption of this ubiquitin–E2 interaction via the UBE2D2 mutation S22R,^39^ results in decreased activity against substrate (**Figure 5G**). We suggest that the ARLD1–ubiquitin interface is necessary for binding, and that site 2 ubiquitin binding constrains ligase activity towards K48 specificity. While the E2 at site 2 cannot be charged in the bound conformation (**Figure S3D**), E2–ubiquitin back-binding interaction is important for activity, presumably by assisting the K48 di-ubiquitin coordination.

## Discussion

HECT ubiquitin ligases commonly feature extended domains distal to the HECT catalytic module that are essential for function. How these domains contribute to substrate recognition, guide chain specificity, and regulate ligase activity is important to understand disease mutations frequently identified in those regions.^4,6,16^ In this study, we determined the structure of the full-length *S. cerevisiae* HECT ubiquitin ligase Tom1 during active ubiquitylation and discovered that the Tom1 N-terminal ARLD repeats coordinate a ubiquitin–E2 architecture which contributes to chain specificity and constrains ligase activity. Tom1 harbors a previously uncharacterized ubiquitin interaction site within this N-terminal domain, stabilizing a distinct conformation of the ligase and hence we refer to it as the “structural” ubiquitin. The repurposing of a functional protein to play structural roles is found in other contexts as well. For example, covalent modification of cullin-RING ligases with ubiquitin-like protein NEDD8 regulates their activity,^43,50^ and actin and actin-related proteins function in the maintenance of higher order chromatin structures such as the SWI/SNF complex.^51,52^ Here, the structural ubiquitin contributes to specificity, which appears to happen at the cost of reduced ligase activity. We hypothesize that in Tom1, the tradeoff in ubiquitin processivity is necessary to confer chain linkage specificity in ubiquitin transfer. A similar molecular mechanism has been reported in *S. cerevisiae* HECT ligase Ufd4, for which structured subunits have been shown to recognize K48 di-ubiquitin chain bound to substrate to direct amplification of K27 poly-ubiquitin chains off the existing di-ubiquitin.^21^ Mutations at this site to abrogate ubiquitin binding slow Ufd4 ligase activity. Combined, these results indicate that structural ubiquitin molecules mediate both ligase activity and chain specificity.

Moreover, our study identified a non-canonical E2 interaction site along the Tom1 scaffold. We hypothesize that the large scale ARLD domain rearrangements act to coordinate the novel UBE2D2–di-ubiquitin recognition interface. Specifically, site 2 ubiquitin interacts non-covalently with the HECT catalytic module as well as with ARLD1 and coordinates site 4 ubiquitin through a K48-linkage. The K48 di-ubiquitin in combination with Tom1 ARLD1/2 forms the non-canonical UBE2D2 interaction site. The E2–ubiquitin interface is stabilized by contacts with a hydrophobic patch on ubiquitin through a literature reported ubiquitin back-binding interaction (**Figure S3D**).^42,46,53^ Similar contacts support the ARLD1–E2 interface as well as HECT–E2 interface on UBE2D2. Superposition of ubiquitin charged onto UBE2D2 on site 2 E2 results in a steric clash with ARLD1/2, suggesting that site 2 UBE2D2 plays a structural role in site 2 K48 di-ubiquitin coordination as opposed to a catalytic role in this conformation.

The molecular architecture of orthologs Tom1 and HUWE1 is highly conserved (**Figures 1B, 1D, S1B, S1D and S1E**) with several HUWE1 patient mutations appearing at conserved interfaces proximal to the ubiquitin and E2 interaction sites we identified (**Figure S4A**).^32,33^ In human cells, HUWE1 activity is constrained: while loss of HUWE1 is not tolerated, patient mutations that result in subtle biochemical changes in HUWE1 activity are associated with diverse pathologies.^16,54,55^ Particularly, two HUWE1 patient mutations (H669Q and G660R) are positioned at conserved residues, which coincide with the identified structural ubiquitin and UBE2D2 binding site (**Figure S4A, S4B**). Adding back HUWE1 domain deletions or patient mutation (H669Q) in a *tom1*Δ background results in a growth deficiency similar to adding back the catalytically inactive form of HUWE1 (**Figure S4B**). Our data for the first-time sheds light on how these distal patient mutations can directly impact the activity of a HECT ubiquitin ligase.

Further mechanistic studies employing chemical probes will be necessary to trap high-energy intermediates and understand the catalytic cycle in the context of the ubiquitin and E2 architecture.^13,19,21^ Though site 1 ubiquitin has been previously identified in the context of the isolated HECT domain,^11,14^ the full-length context of the enzyme was necessary to understand the ubiquitin architecture coordinated proximal to the ligase catalytic cysteine. The precise catalytic geometry and large-scale rearrangements associated with charging of the HECT domain remain to be determined. In conclusion, we combine structural and functional studies to understand the core activity of a HECT ligase implicated in multiple cellular processes and disease pathologies. In the process, we discovered a structural ubiquitin and regulatory mechanism that influences the ligase’s activity and specificity.

## Limitations

While our structural studies into Tom1 engaged in active ubiquitylation captures snapshots of ligase regulation, we do not capture all important intermediates (**Figure S4E**). For example, we do not visualize Tom1 or E2 charged with ubiquitin likely due to the very transient nature of these states. Additional studies of Tom1 in complex with transition state mimetics will be necessary to understand the molecular mechanism underlying how the ubiquitin–E2 architecture guides ligase specificity. Disruption of interfaces contacts influence activity and chain formation on model substrates in our biochemical assays, however, in all classes, we do not directly visualize substrate via cryo-EM.

## Acknowledgments

We thank the staff at the Harvard Cryo-EM Center for Structural Biology for their support during grid screening and data collection, as well as the SBGrid consortium for software support and computing resources.^56^ We thank Milka Kostic for discussion and feedback on the manuscript and all members of the Fischer lab. This study was supported by funding from the Mathers Foundation [Award #64395403]. K.B. is a Damon Runyon Fellow supported by the Damon Runyon Cancer Research Foundation (DRG-2514-24).

## Author Contributions

Conceptualization: K.W., M.H., K.B., D.O. and E.S.F.

Methodology: K.W., M.H., K.B., D.O., A.S., C.J. and S.S.R.B.

Formal Analysis: K.W., M.H., K.B. and A.S.

Investigation: K.W., M.H., K.B. and A.S.

Writing – Original Draft: K.W.

Writing – Reviewing & Editing: S.S.R.B., M.H., K.B. and E.S.F.

Supervision: M.H., S.S.R.B., K.B., K.A.D. and E.S.F.

Project Administration: E.S.F.

Funding Acquisition: E.S.F.

## Declaration of interests

E.S.F. is a founder, scientific advisory board (SAB) member, and equity holder of Civetta Therapeutics, Proximity Therapeutics, Neomorph, Inc. (also board of directors), Stelexis Biosciences, Inc., and CPD4, Inc. (also board of directors). He is an equity holder and SAB member for Avilar Therapeutics, Photys Therapeutics, and Ajax Therapeutics and an equity holder in Lighthorse Therapeutics and Anvia Therapeutics, Inc.. E.S.F. is a consultant to Novartis, EcoR1 capital, Odyssey and Deerfield. The Fischer lab receives or has received research funding from Deerfield, Novartis, Ajax, Interline, Bayer and Astellas. K.A.D. receives or has received consulting fees from Neomorph, Inc and Kronos Bio.

## STAR Methods

### RESOURCE AVAILABILITY

#### Lead contact

Further information and requests for resources and reagents should be directed to and will be fulfilled by the lead contact, Eric S. Fischer.

#### Materials availability

Reagents generated in this study are available from the lead contact with a completed Material Transfer Agreement.

#### Data and code availability

- Cryo-EM maps have been deposited in the EMDB under the following accession codes:

`o` EMD-47058 (apo-Tom1, BS3-crosslinked)
`o` EMD-47035 (apo-Tom1, BS3-crosslinked, HECT focus map)
`o` EMD-47045 (apo-Tom1, BS3-crosslinked, ARLD focus map)
`o` EMD-47057 (Tom1–E2–ubiquitin)
`o` EMD-47046 (Tom1–E2–ubiquitin, focus map)
`o` EMD-47048 (apo-Tom1 closed conformation, non-crosslinked)
`o` EMD-47047 (apo-Tom1 open conformation, non-crosslinked)
`o` EMD-47050 (open class, Tom1 ubiquitylation)
`o` EMD-47049 (closed class, Tom1 ubiquitylation)
`o` EMD-47056 (class 1, Tom1–ubiquitin)
`o` EMD-47055 (class 2, Tom1–ubiquitin)
`o` EMD-47051 (class 3, Tom1–E2–ubiquitin)
`o` EMD-47053 (class 4, Tom1–E2–ubiquitin)
`o` EMD-47052 (class 5, Tom1–E2–ubiquitin)
`o` EMD-47054 (class 6, Tom1–E2–ubiquitin)
- Cryo-EM models have been deposited in the PDB under the following accession codes: 9DNT (apo-Tom1, closed) and 9DNS (Tom1–E2–ubiquitin).
- Uncropped gels and original Western blot images are available through Mendeley Data and are publicly available as of the date of publication. The DOI is listed in the key resources table.
- Any additional information required to reanalyze the data reported in this paper is available from the lead contact upon request.
- This paper does not report original code.

### EXPERIMENTAL MODEL AND STUDY PARTICIPANT DETAILS

#### Yeast strains, growth conditions, and transformation

All yeast strains used in this study (listed in the Key Resources table) were derived from BY4741 (MATa his3Δ1 leu2Δ0 met15Δ0 ura3Δ0). All transformants were verified by auxotrophic selection. Yeast was grown at 30°C in YPD or appropriate synthetic completed (SC) drop-out media. Tom1 and mutants were cloned into pAG425GPD-ccdB (Addgene #14154) and expressed under the GAP promoter. Experiments were initiated with cells at mid-log phase growth as reported in Sung et al. 2016. Yeast transformation was performed by lithium acetate method.^61^

#### Insect cell culture

*Trichoplusia ni* (High Five; Gibco) cells and *Spodoptera frugiperda* (Sf9; Expression Systems) were cultured in SF-4 Baculo Express ICM medium (BioConcept) and ESF 921 medium (Expression Systems), respectively at 27 °C. All cell lines are routinely tested for mycoplasma contamination via PCR.

### METHOD DETAILS

#### Plasmids

The following plasmids were used in this study: “pAG425GPD-ccdB” (GAP. MCS. CYC1. F1 ori. LEU2p. LEU2), Addgene #14154.

#### Antibodies

The following antibodies were used in this study: Immunoblotting: anti-Histone H3 (ABclonal, #A2348, 1:1000), anti-Histone H2B (Abcam, #ab1790, 1:1000), anti-Histone H1 (Abcam, #ab134914, 1:1000), anti-Redd4 (CST, #2516, 1:1000), anti-K48 ubiquitin linkage (CST, #8081, 1:1000), anti-Mcl1 (CST, #5453, 1:1000), anti-mouse 680RD (LI-COR Biosciences, #926-68070, 1:15000), anti-rabbit 800CW (LI-COR Biosciences, #926-32211, 1:15000).

#### *In vitro* ubiquitylation assays

In vitro ubiquitylations reactions were carried out in a total volume of 44 μl with UBA1 at 0.2 μM, UBE2D2 at 0.5 μM, 0.7 μM indicated substrate and Tom1 or HUWE1 at 0.25 μM, buffered in 100 mM NaCl, 25 mM HEPES (pH 7.4), 4 mM ATP and 5 mM MgCl_2_. Either ubiquitin (R&D systems, unlabelled) or recombinant ubiquitin conjugated with fluorescein dye (R&D) at 2.5 μM were added to initiate ubiquitylation. Reactions were allowed to react for the indicated reaction time. Reactions containing Tom1 were conducted at 30°C, while reactions containing HUWE1 were conducted at 37°C.

Reactions containing fluorescent histone or fluorescent ubiquitin were separated by SDS-PAGE and were analyzed by fluorescent imaging on Typhoon FLA 9500 (GE Life Sciences). Reactions without fluorescent histone or ubiquitin were immunoblotted. Samples were separated by SDS-PAGE on either 4-12% Bis-Tris Protein Gels (Thermo Fisher Scientific, #NP0322BOX), 4-20% Mini-PROTEAN TGX Stain-Free Protein Gels (Bio-Rad, #4568096), or 3-8% Tris-Acetate Protein Gels (Thermo Fisher Scientific, #EA03755BOX), and then transferred to a PVDF membrane with an iBlot 2 dry blotting system (Thermo Fisher Scientific, IB21001). Membranes were blocked in Intercept (PBS) Blocking Buffer (LI-COR Biosciences) and incubated with primary antibodies at 4 °C overnight. Following, the membranes were rinsed in Phosphate-buffered saline with Tween 20 (PBS-T), incubated for 1 hour in secondary IRDye-conjugated antibodies (LI-COR Biosciences) and then washed three times in PBS-T prior to imaging on an Odyssey Imaging System (LI-COR Biosciences).

#### Protein expression and purification

UBE2D2 was cloned into pGEX 4T1 derived vectors and expressed as N-terminal GST-TEV fusion proteins in Rosetta (DE3) competent cells (Novagen). UBA1 was cloned into a pLIB derived vector and expressed as N-terminal GST-TEV fusions in High Five cells. Rosetta (DE3) cells were expanded in 1 L of Terrific Broth (TB) containing ampicillin and chloramphenicol at 37 °C shaking at 200 RPM until cell density reached OD_600_ ~0.8. Cells were then induced at 0.2 mM IPTG and grown overnight at 22 °C shaking at 200 RPM, and subsequently harvest by centrifugation (3,300 *g*, 15 min). Wild-type Tom1 and Tom1 mutants were cloned into pAC-derived vectors^62^ and expressed as N-terminal StrepII–TEV fusion proteins in High Five cells. Baculovirus for protein expression was produced by transfection of expression plasmids into Sf9 cells at a density of 0.9 × 10^6^ cells/mL in ESF 921 medium (Expression Systems), following the manufacturer’s instructions. High Five cells were infected at a density of 1.8–2.2 × 10^6^ cells/mL in SF-4 Baculo Express ICM medium (BioConcept) and then harvested 40–60 hours post-infection via centrifugation (1,500 *g*, 20 min).

##### UBA1 purification

Harvested cells were resuspended in lysis buffer [50 mM Tris pH 8.0, 200 mM NaCl, 5 mM TCEP] supplemented with 2 mM PMSF, Aprotinin and Leupeptin and lysed by sonication. The lysate was cleared by high-speed centrifugation (12,000 rpm, 45 min) in Nalgene tubes. The supernatant was incubated with 5 mL GST resin for 1 hour at 4 °C, then spun (2,000 rpm for 10 min) to pellet resin. Resin was resuspended in lysis buffer and transferred to a gravity flow column. Resin was washed in 400 mL of lysis buffer and bound protein was eluted in elution buffer [50 mM Tris pH 8.0, 200 mM NaCl, 5 mM DTT, 20 mM Reduced Glutathione]. Protein was cleaved by TEV protease (1:100) at room temperature for 1 hour, then overnight at 4 °C. Protein was then diluted to 50 mM NaCl in buffer [25 mM HEPES/NaOH pH 7.5, 1 mM TCEP] to bind to Poros 50HQ (Thermo Fisher Scientific), eluting with a linear NaCl-gradient from 20 mM to 1 M. Fractions containing protein were combined, concentrated using centrifugal concentrators, then subject to size exclusion chromatography (Superdex200, Cytiva) in gel filtration buffer [25 mM HEPES/NaOH pH 7.5, 150 mM NaCl, 1 mM TCEP].

##### UBE2D2 purification

Harvested cells were resuspended in lysis buffer [50 mM Tris pH 7.4, 200 mM NaCl, 5 mM DTT] supplemented with 2 mM PMSF and lysed by sonication. The lysate was cleared by ultracentrifugation (25,000 rpm, 30 min). The clarified supernatant was incubated with 5 mL of GST resin for 1 hour at 4 °C, then spun (2,000 rpm for 10 min) to pellet resin. Resin was resuspended in lysis buffer and transferred to a gravity flow column. Resin was washed in 400 mL of lysis buffer and bound protein was eluted in elution buffer [50 mM Tris pH 7.4, 200 mM NaCl, 5 mM DTT, 20 mM Reduced Glutathione]. Protein was cleaved by TEV protease (1:100) at room temperature for 1 hour, then overnight at 4 °C. Protein was then diluted to 20 mM NaCl in buffer [25 mM HEPES/NaOH pH 6.5, 1 mM TCEP] to bind to Poros 50HS (Thermo Fisher Scientific), eluting with a linear NaCl-gradient from 20 mM to 1 M. Fractions containing protein were combined, concentrated using centrifugal concentrators, then subject to size exclusion chromatography (Superdex75, Cytiva) in gel filtration buffer [25 mM HEPES/NaOH pH 7.5, 150 mM NaCl, 1 mM TCEP].

##### Wild-type and mutant Tom1 purification

Harvested cells were resuspended in lysis buffer [50 mM HEPES/NaOH pH 7.2, 350 mM NaCl, 5% (v/v) glycerol, 0.05% (v/v) Tween-20, 2 mM TCEP] supplemented with protease inhibitors and lysed by sonication. The cell lysate was clarified by ultracentrifugation (186,000 *g*, 1 hour), and the clarified supernatant was bound to StrepTactin XT High Capacity resin (IBA) in a gravity flow column. The resin was washed in lysis buffer and protein was eluted in lysis buffer supplemented with 50 mM biotin. Protein was diluted to 300 mM NaCl in buffer [50 mM HEPES/NaOH pH 7.2, 2 mM TCEP] to bind to PorosHQ (Thermo Fishe Scientific), eluting with a linear NaCl-gradient from 225 mM to 750 mM. Fractions containing protein were combined, concentrated using centrifugal concentrators, then subject to size exclusing chromatography (Superose 6 Increase, GE Healthcare) in gel filtration buffer [30 mM HEPES/NaOH pH 7.2, 150 mM NaCl, 2 mM TCEP].

##### BS3-crosslinked Tom1 purification

Harvested cells were resuspended in lysis buffer [50 mM HEPES/NaOH pH 7.2, 350 mM NaCl, 5% (v/v) glycerol, 0.05% (v/v) Tween-20, 2 mM TCEP] supplemented with protease inhibitors and lysed by sonication. The cell lysate was clarified by ultracentrifugation (186,000 *g*, 1 hour), and the clarified supernatant was bound to StrepTactin XT High Capacity resin (IBA) in a gravity flow column. The resin was washed in lysis buffer and protein was eluted in lysis buffer supplemented with 50 mM biotin. Protein was diluted to 300 mM NaCl in buffer [50 mM HEPES/NaOH pH 7.2, 2 mM TCEP] to bind to PorosHQ (Thermo Fishe Scientific), eluting with a linear NaCl-gradient from 225 mM to 750 mM. Fractions containing protein were combined and concentrated using centrifugal concentrators. Tom1 was incubated with 300X molar excess bis(sulfosuccinimidyl)suberate (BS3, Thermo Fisher Scientific) at room temperature for 20 min. Protein samples were quenched in 100 mM Tris pH 7, then subject to size exclusion chromatography (Superose 6 Increase, GE Healthcare) in gel filtration buffer [30 mM HEPES/NaOH pH 7.2, 150 mM NaCl, 2 mM TCEP].

##### Labeling of H2B with BODIPY-maleimide

Site-specific labelling of histone H2B was accomplished by reacting Cys-155 with a maleimide-conjugated fluorophore. Recombinant Histone H2B (Sigma Aldrich, #H2167) was reduced in 10-fold molar excess TCEP at room temperature for 20 minutes, then buffer exchanged using Zeba Spin Desalting Columns (Thermo Fisher Scientific) into labeling buffer [50 mM Tris pH 7.5, 150 mM NaCl, 0.1 mM TCEP] and incubated with 3-fold molar excess BODIPY-maleimide for 3 hours at room temperature, then 4 °C overnight. Excess BODIPY-maleimide was removed by three-rounds of buffer exchange into storage buffer [50 mM Tris pH 7.5, 150 mM NaCl, 0.25 mM TCEP, 10% glycerol] using Zeba Spin Desalting Columns (Thermo Fisher Scientific), and labelling was confirmed by SDS-PAGE and fluorescent imaging on Typhoon FLA 9500 (GE Life Sciences).

#### BS3-crosslinked Tom1 cryo-EM sample preparation and data collection (Data set 1)

R 1.2/1.3 300 grids (Quantifoil) were glow-discharged for 60 s at 15–20 mA and 39 Pa. 8 µL of sample (4.8 mg/mL crosslinked StrepII–TEV–Tom1 and 0.3 mM CHAPSO) were applied twice (4 µL per application) to grids. Grids were vitrified using a Leica EM GP plunge freezer at 90% humidity and 10 °C, with 10 s pre-blot, 3 s blot, 2.7 s post-blot. Grids were imaged with a Thermo Fisher Titan Krios equipped with a Gatan Quantum image filter (20 eV slit width) and a post-GIF Gatan K3 direct electron detector. Screening and data collection were performed using SerialEM (v4.1b). 3 movies (50 frames over 2.8 s, 11,583 movies in total) were acquired per hole (9 holes per stage position), with a total dose of 53.687 e^−^/Å^2^. The defocus ranged from −1.1 to −2.2 µm.

#### Non-crosslinked Tom1 cryo-EM sample preparation and data collection (Data set 2)

Sample was applied twice onto R 1.2/1.3 300 grids (Quantifoil) that were glow-discharged for 60 s at 15– 20 mA and 39 Pa: 4 µL of sample (0.5 mg/mL StrepII–TEV–Tom1 and 0.3 mM CHAPSO) followed by 4 µL of sample (1 mg/mL StrepII–TEV–Tom1 and 0.3 mM CHAPSO). Grids were vitrified using a Leica EM GP plunge freezer at 90% humidity and 10 °C with 10 s pre-blot, 3 s blot, 2.7 s post-blot. Grids were imaged with a Thermo Fisher Titan Krios equipped with a Gatan Quantum image filter (20 eV slit width) and a post-GIF Gatan K3 direct electron detector. Screening and data collection were performed using SerialEM (v4.1b). 2 movies (49 frames over 2.806 s, 4,056 movies in total) were acquired per hole (9 holes per stage position), with a total dose of 55.134 e^−^/Å^2^. The defocus ranged from −0.8 to −2.2 µm.

#### Tom1–E2–ubiquitin (Mg-ATP) cryo-EM sample preparation and data collection (Data set 3)

UBA1 and UBE2D2 were pre-charged for 20 minutes at room temperature in a solution containing 0.5 µM UBA1, 18 µM UBE2D2, 2 mM Mg-ATP (R&D Systems) and 25 µM ubiquitin (R&D Systems), buffered with 1X E3 Ligase Reaction Buffer (R&D Systems). Separately, Tom1 was pre-incubated with histone H2B (Sigma-Aldrich, #H2167) at room temperature at room temperature for 20 minutes at a concentration of 8 µM Tom1 and 16 µM histone H2B. Samples were pre-mixed at a one-to-one volume-volume ratio for 45 seconds and 90 seconds, before addition of detergent and immediate sample application (final sample concentration: 4 µM Tom1, 8 µM histone H2B, 0.25 µM UBA1, 9 µM UBE2D2, 1 mM Mg-ATP, 12.5 µM ubiquitin, 0.2 mM CHAPSO). 4 µL of sample were applied twice to R 1.2/1.3 300 grids (Quantifoil) that were glow-discharged for 60 s at 15–20 mA and 39 Pa. Grids were vitrified using a Leica EM GP plunge freezer at 90% humidity and 10 °C with 10 s pre-blot, 3 s blot, 2.7 s post-blot. Grids were imaged in a Thermo Fisher Talos Arctica equipped with a Gatan K3 direct electron detector. Screening and data collection were performed using SerialEM (v4.1b). 1 movie (50 frames over 4.993 s, 6,604 movies in total) was acquired per hole (9 holes per stage position), with a total dose of 52.55 e^−^/Å^2^. The defocus ranged from −0.8 to −2.2 µm.

#### Tom1–E2–ubiquitin cryo-EM sample preparation and data collection (Data set 4)

UBA1 and UBE2D2 were pre-charged for 20 minutes at room temperature in a solution containing 0.5 µM UBA1, 85 µM UBE2D2, 5 mM Mg-ATP (R&D Systems) and 150 µM ubiquitin (R&D Systems), buffered with 1X E3 Ligase Reaction Buffer (R&D Systems). The pre-charge mixture was then buffer exchanged using Zeba Spin Desalting Columns into gel filtration buffer [30 mM HEPES/NaOH pH 7.2, 200 mM NaCl, 0.5 mM TCEP] to remove excess MgCl_2_. Separately, Tom1 was pre-incubated with histone H2B (Sigma-Aldrich, #H2167) at room temperature at room temperature for twenty minutes at a concentration of 10 µM Tom1 and 20 µM histone H2B. Samples were pre-mixed at a one-to-one volume-volume ratio for 45 seconds and 90 seconds, before addition of detergent and immediate sample application (final sample concentration: 5 µM Tom1, 10 µM histone H2B, 0.25 µM UBA1, 44 µM UBE2D2, 75 µM ubiquitin, 0.2 mM CHAPSO).

R 1.2/1.3 300 grids (Quantifoil) were glow-discharged for 60 s at 15–20 mA and 39 Pa. 8 µL of sample were applied twice to grids (4 µL per application). Grids were vitrified using a Leica EM GP plunge freezer operated at 90% humidity and 10 °C with 10 s pre-blot, 3 s blot, 2.7 s post-blot. Grids were imaged with a Thermo Fisher Titan Krios equipped with a Gatan Quantum image filter (20 eV slit width) and a post-GIF Gatan K3 direct electron detector. Screening and data collection were performed using SerialEM (v4.1b). 3 movies (58 frames over 2.896 s, 18,959 movies in total) were acquired per hole (9 holes per stage position), with a total dose of 56.134 e^−^/Å^2^. The defocus ranged from −0.8 to −2.2.

#### BS3-crosslinked Tom1 cryo-EM data processing (Data set 1)

11,583 movies were corrected for beam-induced motion and contrast transfer function (CTF) was estimated using cryoSPARC Live (Figure S2D).^63^ Low-quality movies were excluded, and 4,216,890 particles were picked using templates from on-the-fly 2D classification. Particles were extracted at 1.32 Å/pixel and cleaned up by two rounds of 2D classification. Ab-initio reconstruction revealed one class of particles corresponding to closed-ring Tom1. The particles associated with this class were further refined by homogeneous refinement, then classified by heterogeneous refinement using one good reference volume and four decoy volumes. Particles from the reference class were retained and fed into six additional iterations of heterogeneous refinement. The resulting 452,490 particles were further polished with reference-based motion correction and CTF-refinement. Duplicate particles with centers closer than 50 Å were removed. This led to a consensus reconstruction at 3.01 Å (EMD-47058) (Figure S2E). For local refinement of N-terminal ARLD1, a soft mask was applied and the signal outside ARLD1–ARLD4 was subtracted. Local refinement led to a reconstruction at 2.96 Å (EMD-47045) (Figure S2F). For local refinement, of the HECT domain, a soft mask was applied and signal external to the HECT domain was subtracted. The particles were then subject to masked 3D classification and particles from six of the twenty classes were subject to local refinement leading to a reconstruction at 3.23 Å (EMD-47035) with 174,094 particles (Figure S2G). The sharpened and unsharpened maps from cryoSPARC, as well as a map post-processed with DeepEMhancer^64^, were used for model building. Tom1 was predicted by AlphaFold2 in 700-1000 amino acid stretches, and was rigid-body fit into the density in ChimeraX^40^, then relaxed into the density by ISOLDE (v1.6.0)^41^ and Rosetta (v3.13).^42^ The resulting model was manually adjusted and built in COOT (v0.9.8.0)^43^ and prepared for refinement with phenix.ready_set (v1.19.2-4158),^44^ followed by refinement using target restraints in phenix.real_space_refine (v1.19.2-4158).^45^ The final model was deposited in the PDB as 9DNT (Figure S2S). Data collection parameters and refinement statistics are available in **Table S2**. Structural biology applications used in this project were compiled and configured by SBGrid.^46^

#### Tom1 cryo-EM data processing (Data set 2)

4,056 movies were corrected for beam-induced motion and contrast transfer function (CTF) was estimated in cryoSPARC (Figure S2A). Low-quality movies were rejected, and 566,274 particles were picked using Topaz (v0.2.5)^47^ trained on selected 2D classes (Figure S7B). Particles were extracted at 1.25 Å/pixel and cleaned up by three rounds of 2D classification. Both 2D classification and ab-initio reconstruction revealed an open and closed conformation of Tom1. Reference volumes representing the two Tom1 classes and four decoy volumes were then used as input volumes in heterogeneous refinement with cleaned-up particles from 2D classification. Particles pertaining to either the Tom1 open or Tom1 closed class were fed each into another round of heterogeneous refinement, respectively. Duplicate particles defined as those centers closer than 50 Å were removed in the Tom1 open and Tom1 closed class, and each class was further refined by homogeneous refinement. The resulting 88,291 particles led to a 3.8 Å map corresponding to the Tom1 closed class (EMD-47048) (Figure S2B), and the resulting 63,906 particles led to a 4.1 Å map corresponding to the Tom1 open class (EMD-47047) (Figure S2C).

#### Tom1–E2–ubiquitin (Mg-ATP) cryo-EM (Data set 3)

6,502 movies were corrected for beam-induced motion and CTF was estimated on-the-fly within cryoSPARC Live (Figure S2H). Low-quality movies were excluded, resulting in 3,159,343 particles picked using Topaz (v0.2.5)^47^ trained on templates from on-the-fly 2D classification. Particles were extracted at 1.32 Å/pixel and cleaned up by two rounds of heterogeneous refinement using input volumes from open Tom1 (Figure S2C), closed Tom1 (Figure S2B) and Tom1–E2–ubiquitin (Figure S2R), as well as decoy volumes generated from ab-initio reconstruction of junk 2D classes. Particles from the Tom1 open conformation were fed into two iterations of heterogeneous refinement, duplicate particles defined as those with centers closer than 50 Å were removed, then the resulting 109,827 particles were refined to a 4.9 Å reconstruction of the Tom1 open conformation (Figure S2I). Particles pertaining to the unbound Tom1 closed conformation were filtered to remove particles with centers closer than 50 Å, then fed into non-uniform refinement to arrive at 496,436 particles forming a 3.7 Å reconstruction of the Tom1 closed conformation (Figure S2P). The resulting 527,441 particles pertaining to the Tom1–E2–ubiquitin class were fed into non-uniform refinement then 3D classification with 10 classes. For each class, duplicate particles defined as those with centers closer than 50 Å were removed, and resulting particles were subject to non-uniform refinement resulting in six classifications at respectively 4.2 Å (52,832 particles, Figure S2J), 4.2 Å (50,872 particles, Figure S2K), 4.1 Å (50,830 particles, Figure S2L), 4.0 Å (52,248 particles, Figure S2M), 4.2 Å (49,830 particles, Figure S2N) and 4.0 Å (49,235 particles, Figure S2O).

#### Tom1–E2–ubiquitin cryo-EM data processing (Data set 4)

18,249 movies were corrected for beam-induced motion and CTF was estimated on-the-fly cryoSPARC Live (Figure S2Q). Low-quality movies were excluded, and 11,388,836 particles were picked using Topaz (v0.2.5)^47^ trained on selected 2D classes. Particles were extracted at 1.62 Å/pixel then cleaned by four iterations of heterogenous refinement with input volumes [Tom1 open (Figure S2C), Tom1 closed (Figure S2B) and Tom1–E2–ubiquitin (Figure S2R)] and decoy volumes. The resulting 4,391,455 particles in either the Tom1 closed or Tom1–E2–ubiquitin class were fed into another iteration of heterogenous refinement with three junk classes. The 1,211,061 particles in the Tom–E2–ubiquitin class were refined by non-uniform refinement, forming a 3.6 Å reconstruction. The resulting particles were then fed into 3D classification. Particles from four of the ten classes were fed into heterogeneous refinement, then non-uniform refinement forming a 3.3 Å reconstruction with 247,456 particles. Particles were then re-extracted at 1.1 Å/pixel, duplicate particles defined as those with centers closer than 50 Å were removed and polished by per-particle CTF estimation and reference-based motion correction. The resulting 246,341 particles were fed into non-uniform refinement forming a ~2.8 Å reconstruction of the Tom1–E2–ubiquitin complex. A local mask was applied to arrive at a focused refinement (EMD-47046). The sharpened and unsharpened maps from cryoSPARC, in addition to a map post-processed with DeepEMhancer, were used for model building. Domains from the apo-Tom1 model as well as UBE2D2 and ubiquitin (PDB: 4V3L) were rigid-body fit into the final map in ChimeraX.^40^ Models were relaxed into the density using a combination of ISOLDE (v1.6.0)^41^ and Rosetta (v3.13).^42^ The resulting model was manually adjusted and built in COOT (v0.9.8.0)^43^ and prepared for refinement with phenix.ready_set (v1.21-5207) and phenix.real_space_refine (v1.21-5207)^45^. The final model was deposited in the PDB as 9DNS and all maps were deposited in the EMDB under EMD-47057 (Figure S2T). Data collection parameters and refinement statistics are available in **Table S1**.

#### LC-MS analysis for ubiquitin linkage analysis

*In vitro* ubiquitylation reactions were carried out in a total volume of 60 μL of 100 mM NaCl, 25 mM HEPES (pH 7.4), 4 mM ATP and 5 mM MgCl_2_ with 0.2 μM UBA1, 0.5 μM UBE2D2, 0.7 μM histone H2B and 0.25 μM Tom1 WT or mutant. Either ubiquitin (R&D systems, unlabeled) or recombinant ubiquitin conjugated with fluorescein dye (R&D) were added to initiate ubiquitylation at 2.5 μM. Reactions proceeded for 90 minutes at 30°C and were quenched to 2M urea. Substrate ubiquitylation was confirmed by immunoblot.

Proteins were denatured in 8 M urea in HEPES pH 7.8 for 15 minutes. Disulfides were reduced in 10 mM TCEP for 30 minutes and alkylated in 10 mM iodoacetamide for 45 minutes. Alkylation was quenched with a 10 minute incubation in 10 mM DTT. Urea was diluted to 1 M with 50 mM HEPES pH 7.8 prior to digestion. Proteins were first digested with LysC (1:50; enzyme:protein) for 16 hours at 37°C, then digested in trypsin (1:50; enzyme:protein) for 6 hours at 37°C. Samples were acidified with formic acid to a pH of ~2-3. Stable isotope-labeled peptide standards were spiked in (Cell Signaling Technologies): 1 pmol ubiquitin aa 43-54 (LIFAG**<u>K</u>**Q**<u>L</u>**EDGR, Lys-GlyGly [K6], heavy Leu [L8]) and 1 pmol ubiquitin aa 64-72 (ESTLHLV**<u>L</u>**R, heavy Leu [L8]). Samples were desalted using C18 solid phase extraction plates (SOLAμ, Thermo Fisher Scientific). Desalted peptides were dried in a vacuum-centrifuged and reconstituted in 0.1 % formic acid.

Digested peptides were analyzed on an Orbitrap Eclipse mass spectrometer coupled to an Ultimate 3000 RSLCnano (Thermo Fischer Scientific). Peptides were separated over a 50-cm C18 column (ES803A, Thermo Fischer Scientific) with a 70-min gradient of 6-30% acetonitrile in 0.1% formic acid and electrosprayed (1.9 kV, 300°C) with an EasySpray ion source (Thermo Fischer Scientific). Precursor ion scans (375-1,325 m/z) were obtained in the orbitrap (120,000 resolution, profile). Data dependent MS^2^ scans (n = 2 in 15 s, exclusion duration = 30 s) were acquired in the orbitrap following HCD fragmentation (35% NCE, 0.7 m/z isolation, 30,000 resolution).

Raw data were searched against a database containing the reaction component sequences and common contaminant proteins using SEQUEST in Proteome Discoverer 2.4, permitting a mass tolerance of ±10 ppm, 2 missed cleavages by trypsin, static carbamidomethylation of Cys, and the following variable modifications: oxidation of Met, phosphorylation of Ser/Thr/Tyr, ubiquitylation of Lys, and heavy Val, Pro, Leu, Iso. Peptide spectral matches were validated using a target/decoy approach (1% FDR). Precursor ions were quantified with the Minora Feature Detector, Feature Mapper, and Precursor Ion Quantifier nodes, and relative changes in ubiquitin linkages were quantified against ub-AQUA peptide standards.

## Supplemental Figure Titles and Legends

**Figure S1. Characterization of Tom1 structure and conservation**

**(A)** SDS-PAGE analysis of purified Tom1 pre- and post-BS3 crosslinking. Pooled fractions following size exclusion chromatography (Superose6, Cytiva) indicated.

**(B)** Domain map of human HUWE1 and *Nematodica spERT5* HUWE1_N_, indicating domain boundaries and amino acids stretched by ARLD1-4. Color scheme for domain annotations are kept constant from (Figure 1C).

**(C)** Representative 2D classes of Tom1 pre- and post-crosslink.

**(D) (Left)** Overlay of human HUWE1 (PDB: 7JQ9) and Tom1. Among orthologs, the root-mean square distance (RMSD) between orthologs is 1.4 Å and 2.5 Å for the HECT domain and ARLD1, respectively. **(Right)** HUWE1, Tom1 and HUWE1_N_ HECT domains aligned on ARLD4 to depict mobility of the HECT domain relative to closed-ring architecture.

**(E)** Conservation analysis of Tom1 among 125 orthologs. Consurf^57–60^ conservation score mapped by residue and associated domain annotation.

**Figure S2. Cryo-EM processing and Model Validation of Cryo-EM Datasets 1-4**

**(A)** Processing of non-crosslinked Tom1 (Dataset 2). Summary of cryo-EM processing workflow, from representative raw micrograph (low-pass filtered to 5 Å, scale bar-included) to representative 2D classes, revealing large-scale conformational rearrangements of the ligase. Ab-initio reconstruction resulted in two significant classes associated to Tom1.

**(B)** Overview of processing workflow for closed-ring conformation of Tom1 (Dataset 2). Particles belonging to the colored map were fed into further rounds of heterogeneous and homogeneous refinement, resulting in the final reconstruction at 3.8 Å. Map colored by local resolution is depicted to the left. (Top to bottom): FSC plot for closed-ring Tom1; viewing direction distribution for closed-ring Tom1; 3DFSC plot and directional distribution histogram for closed-ring Tom1.

**(C)** Overview of processing workflow for open conformation of Tom1 (Dataset 2). Particles belonging to the colored map were fed into further rounds of heterogeneous and homogeneous refinement, resulting in the final reconstruction at 4.1 Å. Map colored by local resolution is depicted to the left. (Top to bottom): FSC plot for open Tom1; viewing direction distribution for open Tom1; 3DFSC plot and directional distribution histogram for open Tom1.

**(D)** Overview of processing workflow for BS3-crosslinked Tom1 (Dataset 1), from representative raw micrograph filtered to to 5 Å, scale bar-included. Representative 2D classes of closed-ring Tom1 and junk particle classes. Resulting ab-initio models displayed. Further round of refinement, particle polishing, masked classification and local refinements depicted.

**(E)** (Left to Right, top) The final consensus map (Dataset 1); the final map fed into DeepEMhancer; final map colored by local resolution. (Left to Right, bottom) Viewing directional distribution; FSC plot; 3DFSC plot and directional distribution histogram.

**(F)** (Left to Right, top) Local refinement of ARLD1-ARLD4 interface (Dataset 1); the final map fed into DeepEMhancer; final map colored by local resolution. (Left to Right, bottom) Viewing directional distribution; FSC plot; 3DFSC plot and directional distribution histogram.

**(G)** (Left to Right, top) Local refinement of the HECT domain (Dataset 1); the final map fed into DeepEMhancer; final map colored by local resolution. (Left to Right, bottom) Viewing directional distribution; FSC plot; 3DFSC plot and directional distribution histogram.

**(H)** Overview of processing workflow for Tom1–E2–ubiquitin occupancy classes (Dataset 3), from representative raw micrograph filtered to to 5 Å, scale bar-included.

**(I)** The final map of open conformation Tom1 (Dataset 3). Page 2, left to right: Final map colored by local resolution; FSC plot; 3DFSC plot and directional distribution histogram; viewing directional distribution.

**(J)** The final map of class (i) Tom1–E2–ubiquitin (Dataset 3). Page 2, left to right: Final map colored by local resolution; FSC plot; 3DFSC plot and directional distribution histogram; viewing directional distribution.

**(K)** The final map of class (ii) Tom1–E2–ubiquitin (Dataset 3). Page 2, left to right: Final map colored by local resolution; FSC plot; 3DFSC plot and directional distribution histogram; viewing directional distribution.

**(L)** The final map of class (v) Tom1–E2–ubiquitin (Dataset 3). Page 2, left to right: Final map colored by local resolution; FSC plot; 3DFSC plot and directional distribution histogram; viewing directional distribution.

**(M)** The final map of class (iv) Tom1–E2–ubiquitin (Dataset 3). Page 2, left to right: Final map colored by local resolution; FSC plot; 3DFSC plot and directional distribution histogram; viewing directional distribution.

**(N)** The final map of class (iii) Tom1–E2–ubiquitin (Dataset 3). Page 2, left to right: Final map colored by local resolution, at a contour; FSC plot; 3DFSC plot and directional distribution histogram; viewing directional distribution.

**(O)** The final map of class (vi) Tom1–E2–ubiquitin (Dataset 3). Page 2, left to right: Final map colored by local resolution, at a contour; FSC plot; 3DFSC plot and directional distribution histogram; viewing directional distribution.

**(P)** The final map of closed-ring conformation Tom1 (Dataset 3). Page 2, left to right: Final map colored by local resolution, at a contour; FSC plot; 3DFSC plot and directional distribution histogram; viewing directional distribution.

**(Q)** Overview of processing workflow for Tom1–E2–ubiquitin complex (Dataset 4), from representative raw-micrograph, low pass-filtered to 5 Å. Particles belonging to the colored map were fed into further rounds of refinement, 3D classification and particle polishing. Consensus and focused map are depicted with the following data: final map, final map colored by local-resolution, viewing directional distribution, DeepEMhancer sharpened map, FSC plot and 3DFSC plot and directional distribution histogram.

**(R)** Overview of processing workflow for Tom1–E2–ubiquitin initial volume, collected under identical conditions to Dataset 3 on a Talos Arcitica microscope (Thermo Fisher Scientific).

**(S)** Model-to-Map FSC plot of Tom1-BS3, as in Figure 2.

**(T)** Model-to-Map FSC plot of Tom1–E2–ubiquitin, as in Figure 4.

**(U)** Density supporting site 2 K48 di-ubiquitin linkage. Model of Tom1–E2–ubiquitin complex depicted in the final consensus map resulting from Dataset 4 (Figure S2Q).

**(V)** Density supporting site 1-3 K48 di-ubiquitin linkage. Model of Tom1–E2–ubiquitin complex depicted in the final consensus map resulting from Dataset 4 (Figure S2Q).

**Figure S3. Structural analysis of Tom1–E2–ubiquitin complex**

**(A)** Structural superposition of Tom1 HECT–ubiquitin site 1 to reported NEDD4– ubiquitin complexes (left: PDB 4BBN and right: PDB 2XBB). These structures suggest the N-lobe ubiquitin binding site is conserved among HECT family members. Structures are aligned by the N-lobe of each respective HECT domain. Ubiquitin RMSD is reported.

**(B)** Overlay of HECT domain from apo-Tom1 (white) and Tom1–E2–ubiquitin complex (salmon) in complex with site 1 ubiquitin (light yellow) and site 2 ubiquitin (dark yellow) indicates conformational rearrangement upon ubiquitin binding. For clarity, the catalytic cysteine is highlighted in red. Complexes are aligned on ARLD4 (aa 2700-2889), displayed in light gray.

**(C)** Overlay of apo-Tom1 (HECT domain pink, N-terminal domains gray) and Tom1 in Tom1–E2–ubiquitin complex (white) indicates the conformational rearrangements of the closed-ring architecture upon ubiquitin–E2 binding.

**(D)** Alignment of UBE2D2–ubiquitin back-binding interaction (site 2 UBE2D2 green and site 2 di-ubiquitin dark yellow) with reported UBE2D2–ubiquitin complex (UBE2D2 light green, ubiquitin brown; PDB: 4V3L), aligned on UBE2D2. Overlay of charged UBE2D2 (PDB: 4V3L) with site 2 UBE2D2 results in a steric clash with the Tom1 ARLD repeats in the bound conformation.

**(E)** Left: Structural alignment of UBE2D2~Ub^D^~HECT^NEDD4L^ (PDB: 3JVZ) with Tom1 HECT N-lobe in the Tom1–E2–ubiquitin complex suggests donor ubiquitin may be transferred to the Tom1 HECT domain in the site 1 E2–HECT bound conformation; Middle: Structural alignment of HECT^NEDD4^~Ub^D^ (PDB: 4BBN) with the Tom1 N-lobe in the Tom1–E2–ubiquitin complex; Right: Structural alignment of HUWE1~Ub^D^ (PDB: 6XZ1) with the Tom1 N-lobe in the Tom1–E2–ubiquitin complex indicates a steric clash with ARLD1.

**Figure S4. Structural and functional characterization of Tom1/HUWE1 conservation**

**(A)** Left: HUWE1 (PDB: 7JQ9) aligned by domain to Tom1–E2–ubiquitin complex. Surface representation of HUWE1 patient mutations colored in fuchsia. Right: Surface representation of Tom1 in Tom1–E2–Ubiquitin complex, with residues colored by conservation score as in (Figure 1E).

**(B)** Reported HUWE1 patient mutations which coincide with E2 and ubiquitin interfaces along Tom1.^16^

**(C)** Cells of the indicated genotypes are spot platted on YPD and incubated at 30°C or 37°C for three days.

