## Supplementary figures and images for "Structural ubiquitin contributes to K48 linkage-specificity of the HECT ligase Tom1"

### Figure S1

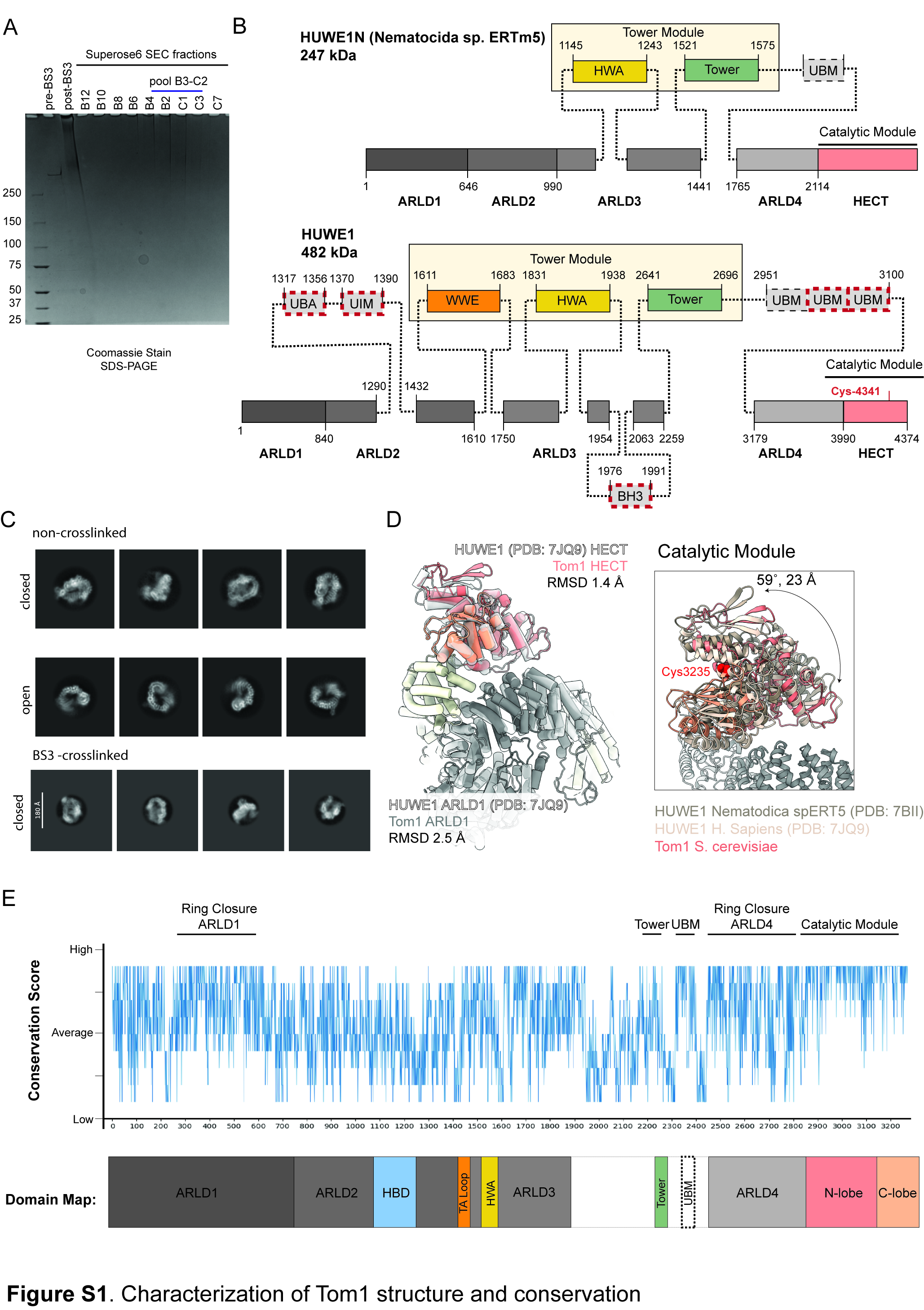

### Figure S2

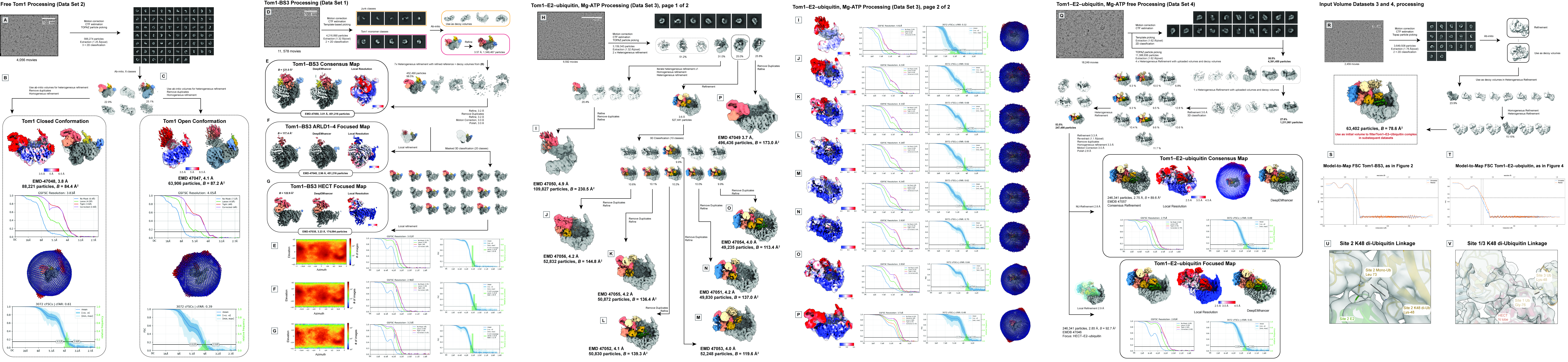

### Figure S3

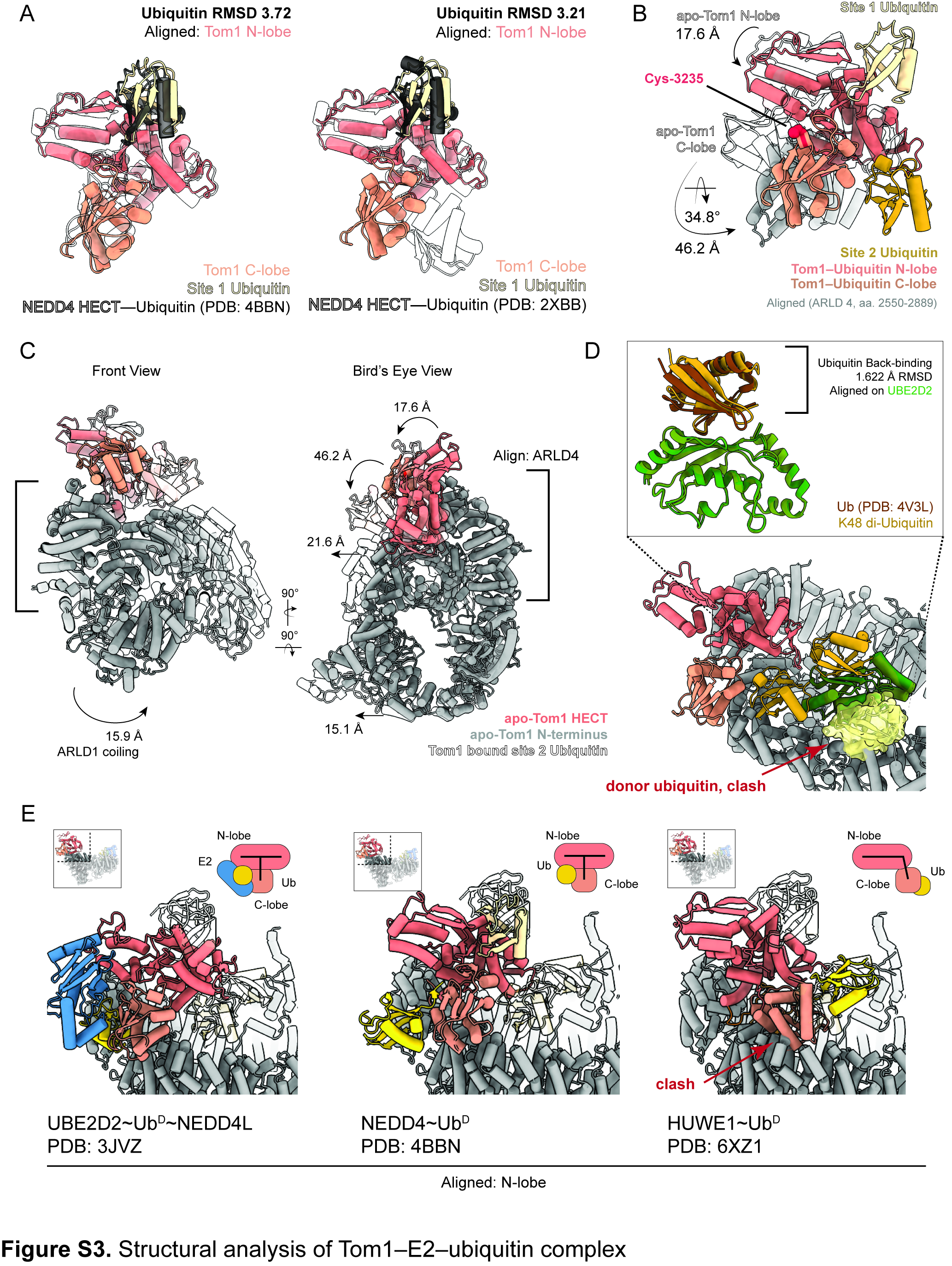

### Figure S4

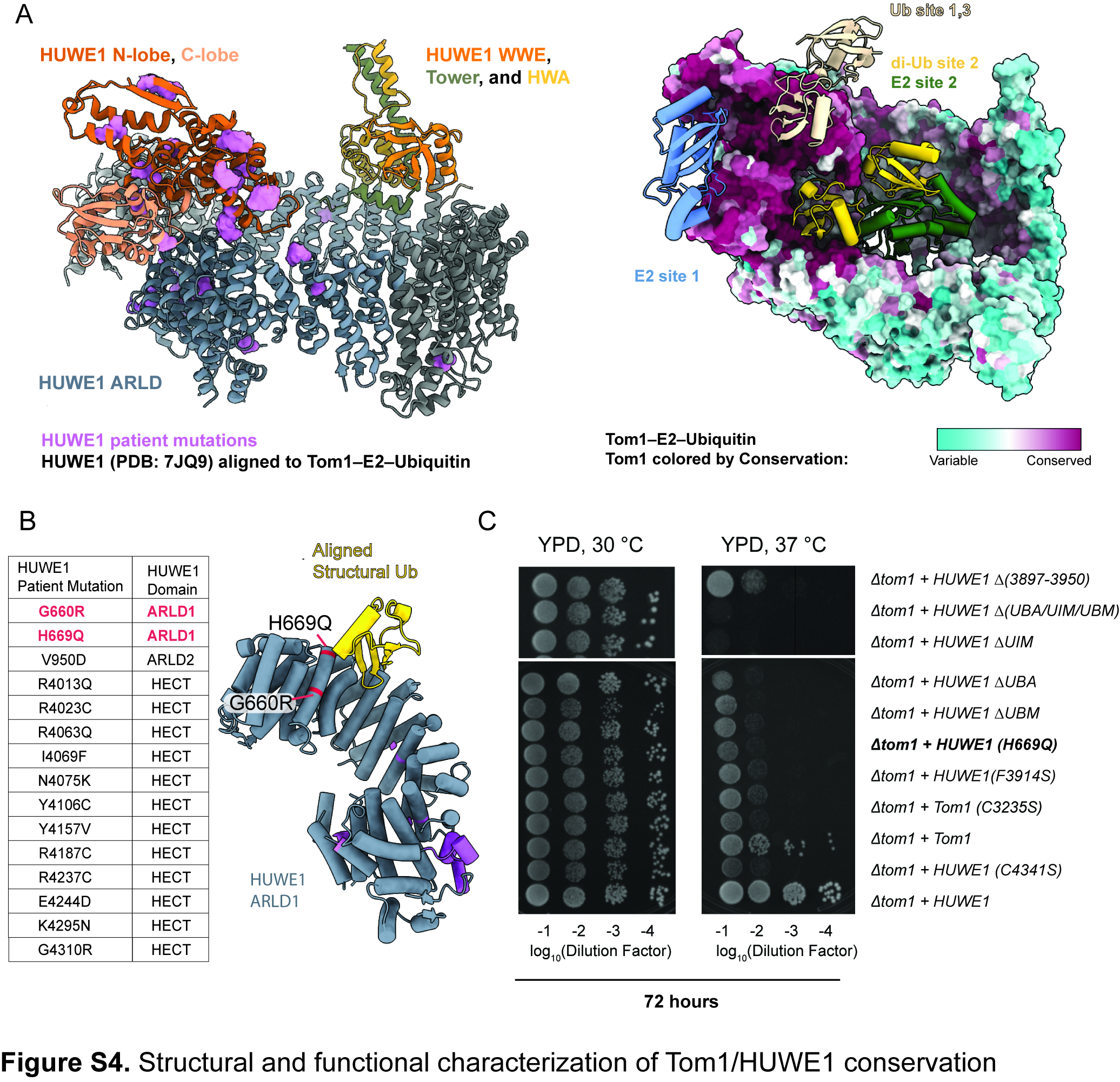
